# Diversity and dynamics of the CRISPR-Cas systems associated with *Bacteroides fragilis* in human population

**DOI:** 10.1101/2021.09.09.459629

**Authors:** Tony J. Lam, Kate Mortensen, Yuzhen Ye

## Abstract

CRISPR-Cas systems are adaptive immune systems commonly found in prokaryotes that provide sequence-specific defense against invading mobile genetic elements (MGEs). The memory of these immunological encounters are stored in CRISPR arrays, where spacer sequences record the identity and history of past invaders. Analyzing such CRISPR arrays provide insights into the dynamics of CRISPR-Cas systems and the adaptation of their host bacteria to rapidly changing environments such as the human gut. In this study, we utilized 601 *Bacteroides fragilis* genome isolates from 12 healthy individuals, 6 of which include longitudinal observations, and 222 available *B. fragilis* reference genomes to update the understanding of *B. fragilis* CRISPR-Cas dynamics and their differential activities. Analysis of longitudinal genomic data showed that some CRISPR array structures remained relatively stable over time whereas others involved radical spacer acquisition during some periods, and diverse CRISPR arrays (associated with multiple isolates) co-existed in the same individuals with some persisted over time. Furthermore, features of CRISPR adaptation, evolution, and microdynamics were highlighted through an analysis of host-MGE network, such as modules of multiple MGEs and hosts, reflecting complex interactions between *B. fragilis* and its invaders mediated through the CRISPR-Cas systems. This work demonstrates the power of using culture-based population genomics to reveal the activities and evolution of the CRISPR-Cas systems associated with gut bacteria in human population. We made available of all annotated CRISPR-Cas systems and their target MGEs, and their interaction network as a web resource at https://omics.informatics.indiana.edu/CRISPRone/Bfragilis.

## Introduction

Microorganisms play a crucial role in human health by forming endosymbiotic relationships with their hosts and other microorganisms. These complex networks of microbial communities found throughout various environments, particularly in the human gut, are referred to as microbiomes [1, 2, 3]. Aside from bacteria-host interactions, bacteria are constantly engaged in an evolutionary arms-race with mobile genetic elements (MGEs), such as phage and plasmids. To defend against antagonistic actors, prokaryotes have developed a variety of mechanisms to alleviate such threats, one of which are CRISPR-Cas systems, an adaptive immune system that provides sequence-specific defense against invading MGEs [4, 5, 6].

CRISPR-Cas systems are highly prevalent, existing in approximately half of bacterial and most of the archaeal genera [7, 8, 9, 10, 11]. The extreme diversity of CRISPR-Cas systems is reflected by their ever-changing classification scheme, owing to the constant discovery of new CRISPR-Cas system types and subtypes [5, 12, 13]. CRISPR-Cas systems can be grouped into two main classes: Class I and Class II CRISPR-Cas Systems. Class I CRISPR-Cas Systems includes Types I, II and V and use a complex of Cas proteins to degrade foreign nucleic acids. Class II CRISPR-Cas Systems include Types II, V, and VI and use a single, large Cas protein for the same purpose (Type II, V and VI use Cas9, Cas12 and Cas13, respectively) [14]. The diversity of CRISPR-Cas systems provides a fitness edge against invaders and is suggested to be a product of advantageous evolution [15, 16, 17]. Similarly, evolution of invaders have been observed to occur in tandem with host adaptive immunity as to evade host defense mechanisms, such as anti-CRISPR genes [4, 5, 18, 19, 20, 21].

CRISPR arrays are comprised of short DNA segments, known as spacers, and these provide a cornerstone to CRISPR-Cas derived adaptive immunity. Spacers, which are originally segments of invaders’ genomes, retain the memory of past immunological encounters, and are primarily acquired as a result of Cas protein complex mediated acquisition [5]. Newly acquired spacers are typically integrated towards the leader ends of arrays [22, 23]. Additionally, leader sequences usually found upstream of CRISPR arrays are attributed to the efficiency of CRISPR-Cas derived immune response [24]. Several studies have also suggested that spacer acquisition remains possible through several alternative means such as homologous recombination [22, 25, 26], and ectopic spacer integration where spacers are inserted into the middle of arrays as a result of leader sequence mutations [24, 27]. While CRISPRs hold immunological memory of past encounters with some arrays spanning several hundred spacers long [28], CRISPR arrays are typically found to be on average less than 30 spacers long suggesting that some spacers are purged over time [29]. While a specific underlying mechanism of CRISPR array maintenance has not yet been found, various studies have suggested several mechanisms of spacer loss, such as spontaneous deletions, recombination, and DNA polymerase slippage during replication [25, 30, 31, 32].

In recent years, much effort has been placed into better understanding the interactions of microbiomes and their host, as well as, the potential modulation of the human microbiome to improve human health. One particular member of the microbiome, *B. fragilis*, has been proposed as a potential probiotic due to its ability to facilitate the alleviation of certain disease conditions [33]. In contrast, Bacteroidetes is one of the most common genera of bacteria in the lower intestinal tract, and while this member of the microbiome only accounts for a small fraction (∼ 2%) of the total Bacteroides found in the gut microbiome, this species contributes to over 70% of Bacteroides infections [34, 35, 36]. This is due to *B. fragilis*’ extensive pan-genome and susceptibility to horizontal gene transfer events. As a result, certain strains of *B. fragilis* have become a known pathobionts and opportunistic pathogens [37, 38, 39]. The perplexing interplay between the pathogenic and probiotic nature of *B. fragilis* strains highlights the importance of understanding pathobiont evolutionary dynamics, elements that contribute to a species’ pathogenicity, and CRISPR-Cas dynamics. Many studies of the adaptation process of the CRISPR-Cas system involved an individual bacterial species challenged with invaders in controlled assays. Taking advantage of the increasing number *B. fragilis* reference genomes, and more importantly, the large number of *B. fragilis* isolates from 12 individuals, we re-investigated the CRISPR-Cas systems in *B. fragilis* in its natural living environment. The availability of hundreds of time-resolved genomes from *B. fragilis* isolates from 6 individuals (some involving multiple time points) allowed us to investigate both the intra- and inter-personal dynamics of interactions between *B. fragilis* and their invaders, and expand upon previous surveys of *B. fragilis* CRISPR-Cas systems [39]. Insights into how *B. fragilis* interacts with its invaders, as well as its CRISPR-Cas systems confer immunity help provide insights into factors that contribute to *B. fragilis* virulence, horizontal gene transfer, and evolution.

## Methods

### Genomic data processing and assembly

Reads from 601 *B. fragilis* isolates from the Zhao et al. study [40] were downloaded from the NCBI BioProject Accession PRJNA524913, henceforth referred to as the ‘Zhao2019 dataset’. Raw shotgun sequencing reads were trimmed using Trimmomatic v0.39 [41] (parameters used: LEADING:5 TRAILING:5 SLIDINGWINDOW:4:10 MINLEN:60). Trimmed reads were then assembled using SPAdes v3.12 [42] with default settings. FragGeneScan [43] was then used to predict protein coding genes of metagenome assemblies.

A total of 222 *B. fragilis* reference genomes, 16 complete and 202 draft genomes, were downloaded from the NCBI ftp website as of Jan 18, 2021. A list of genomes included in this analysis can be found at the CRISPRone website http://omics.informatics.indiana.edu/CRISPRone/Bfragilis.

### Characterization of CRISPR-Cas systems

To identify CRISPR-Cas systems in *B. fragilis* genomes, we utilized CRISPRone [44] which predicts both CRISPR arrays and *cas* genes within a given input genome sequence. Predicted CRISPR-Cas systems were then further refined through a reference based approach. Repeat sequences of CRISPRone predicted spacers were extracted and clustered to obtain consensus reference repeats using CD-HIT-EST [45] with 85% sequence identity. Consensus reference repeats were then used as input for CRISPRAlign [46], a reference based approach to identify CRISPR arrays. As the exact boundaries of CRISPR arrays predicted by *de novo* approaches may sometimes be blurred due to small CRISPR arrays, sequencing errors, and mutations in repeat sequences, we utilize a reference based approach to redefine the repeat-spacer boundaries of CRISPR arrays predicted by CRISPRone.

To compare spacer sequences across different arrays, reduce spacer redundancy, and the eventual computation of spacer content heterogeneity, spacers were clustered with CD-HIT-EST [45] at 85% sequence identity. An 85% sequence identity was used to provide greater flexibility in spacer sequences, and allow for a small amount of sequence variation either due to sequencing error or real mutations found between individual spacers. Spacer sequences that clustered together were considered identical spacer sequences. Spacer clusters were reserved for downstream computation of spacer content heterogeneity (Figure 1A) and construction of compressed spacer graphs (Figure 1B).

**Figure 1:**
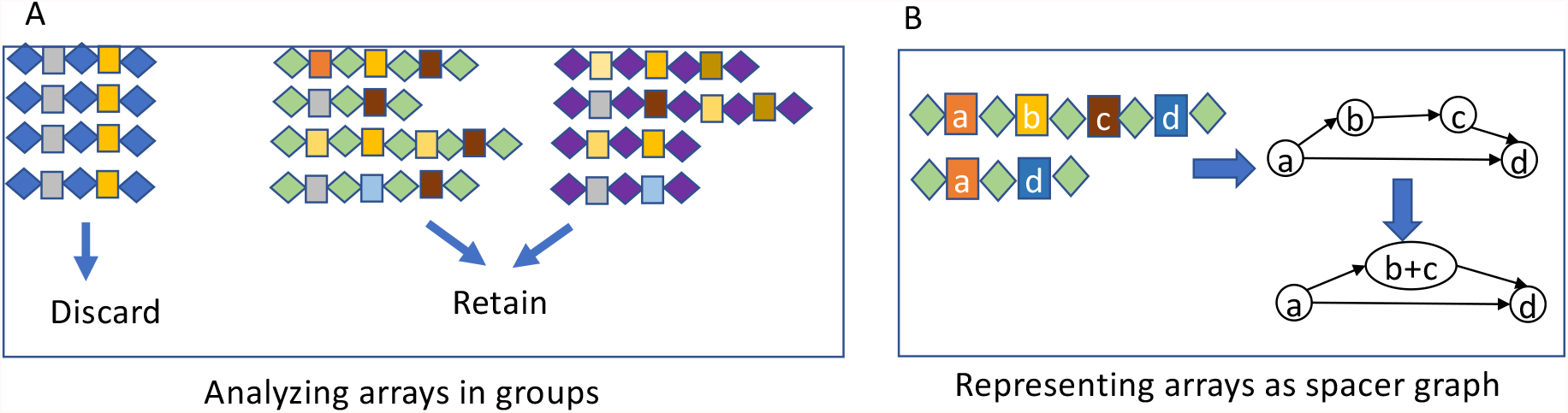
Approaches used for the identification and refinement of the CRISPR arrays and construction of spacer graphs. (A) CRISPR arrays are analyzed in groups such that each group shares identical or very similar repeats (repeats are shown as diamonds and spacers are shown as boxes). CRISPR arrays containing no spacer variance or no CRISPR spacer hetergeneity are discarded. (B) Example of spacer sharing CRISPR arrays can be represented as a simplified graphical structure (spacer graph), in which the edges record the ordering of the spacers in arrays.

In some cases, it may be difficult to differentiate between true CRISPR-Cas systems and false positive CRISPR-Cas systems (e.g., false CRISPR-arrays, inactive CRISPR-Cas systems). While manual curation can help filter out some of these issues, it becomes difficult to screen out hundreds to thousands of genomes. To help filter out false positive arrays and inactive CRISPR-Cas systems, we propose a metric of heterogeneity to measure the rate of change (i.e., growth and turnover of spacers) in CRISPR arrays with the assumption that CRISPR arrays of active CRISPR-Cas systems undergo active expansion and turnover of spacers. Here we define spacer content heterogeneity score as:

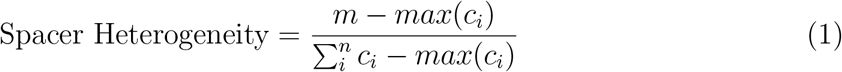

Where *n* is defined as the number CRISPR arrays, with each CRISPR array containing *c*_1_, *c*_2_,…, *c*_*n*_ unique spacers (in some rare cases, CRISPR arrays may contain multiple copies of the same spacer, which will be considered as one spacer) and *m* denotes the number of unique spacers found from all *n* arrays combined. Spacer heterogeneity scores range from 0 to 1, where 0 indicates no spacer heterogeneity (i.e., two CRISPR arrays share all spacers), and 1 indicates the greatest possible extent of spacer content heterogeneity (i.e. two CRISPR arrays share no spacers).

Because spacer content heterogeneity alone is not enough to rule out false positive or inactive CRISPR-Cas systems, predicted CRISPR-Cas systems were further filtered out by coupling spacer content heterogeneity with gene content information. CRISPR groups that lack spacer content heterogeneity and had no adjacent *cas* genes were considered inactive or false positive, and thus discarded from further analysis; all filtered arrays were also manually inspected prior to their removal.

### Compressed spacer graph for summarizing the sharing of spacers among a group of CRISPR arrays

Compressed spacer graphs [47] were constructed for each CRISPR-Cas type to summarize and illustrate CRISPR array dynamics. For every spacer in a given array, where each spacer was represented by a node of its representative spacer cluster, a directed edge was built between nodes of neighboring spacers in sequential order. Once all CRISPR arrays were represented in the graph structure, the spacer graph was then simplified by collapsing neighboring nodes if two neighboring nodes shared an “in-degree” and “out-degree” equal to or less than one (Figure 1B). Compressed spacer graphs highlight CRISPR array structure and dynamics (e.g. branching structures representing spacer gain and loss). Arrays that share no spacers result in disconnected components in the compressed spacer graph.

### Mobile Genetic Element Databases

A collection of mobile genetic element (MGE) databases were gathered, including phage and plasmid databases. The phage databases included the Gut Phage Database [48] (GPD), MicrobeVersusPhage [49] (MVP) database, and the reference viral database [50] (RVDB). The plasmid databases included the Comprehensive and Complete Plasmid Database [51] (COMPASS), and PLSDB [52]. The phage and plasmid databases included sequences from the NCBI reference database, NCBI nucleotide database, MGEs identified from metagenomic assemblies, and prophages identified in prokaryotic genomes. We collectively refer to these databases as the ‘MGE database’ for simplicity.

### Identification of CRISPR Targets

All unique spacer sequences extracted from *B. fragilis*’ CRISPR arrays were queried against the MGE database using BLASTN [53] to search for putative invaders that were targeted by *B. fragilis*. For this analysis, we used all unique spacers instead of 85%-similarity nonredundant set to increase the search sensitivity. Results were filtered to retain hits with a greater than 90% sequence identity, query coverage per hsp greater than 80%, and an e-value of less than 0.001. We noticed that even after dereplication by dRep [54] (with default parameters), there was still a large redundancy in the identified MGEs. Instead, we devised a greedy algorithm to select the minimum number of MGEs that collectively contain all protospacers matching the spacers. Similarly, we selected the minimum number of *B. fragilis* isolates that contained all identified spacers and only included them in the network. Selected MGEs and isolates are then used for building spacer-MGE and host-MGE networks. In the spacer-MGE network, spacer sequence clusters (called spacers for simplicity) and MGEs are represented as nodes and an edge is added between a spacer node and MGE node if the MGE contains a segment that matches the spacer (i.e., protospacer). In the host-MGE network, an edge is added to a host and a MGE if the host and MGE pair contain at least one matching protospacer and spacer. For MGEs that are phages (or prophages), we applied PhaGCN [55] to assign their taxonomic groups (ICTV [56] families). All visualizations and manual inspection of the networks were performed using Cytoscape [57].

## Results

### CRISPR-Cas systems in *B. fragilis*

To better understand the dynamics of CRISPR-Cas systems within *B. fragilis*, we analyzed a total of 823 *B. fragilis* genomes, which included 222 NCBI reference genomes and 601 isolates from the Zhao2019 dataset. Our analysis showed that among all *B. fragilis* genomes, three types of CRISPR-Cas systems were identified, Type I-B, Type II-C, and Type III-B. Example illustrations of these CRISPR-Cas systems are depicted in Figure 2A, and Table 1 shows the signature *cas* genes and the repeat sequence of the CRISPR arrays for these CRISPR-Cas systems. We note that besides the three types of predicted CRISPR-Cas systems, additional putative CRISPR arrays were predicted in a *de novo* fashion. However, they were deemed to be false CRISPR arrays due to their lack of spacer content heterogeneity, despite the fact that they superficially contain the repeat-spacer structures (see details of these CRISPR artifacts and reasons why there were discarded at the supplementary website). Among the discarded CRISPR groups include a putative fourth CRISPR-Cas system that was previously reported in *B. fragilis genomes* [39]. This CRISPR-like artifact was found in 137 out of the 222 (61.7%) reference *B. fragilis* genomes, and isolates of *B. fragilis* in 10 out of 12 individuals in the Zhao2019 dataset (other CRISPR artifacts were rare, found in one or very few genomes). Additionally, this CRISPR artifact was predicted to contain protein-coding genes encoding for transcriptional regulators in some genomes (e.g., CP036550.1 and CP0811922.1), further suggesting that it is unlikely a genuine CRISPR.

**Figure 2:**
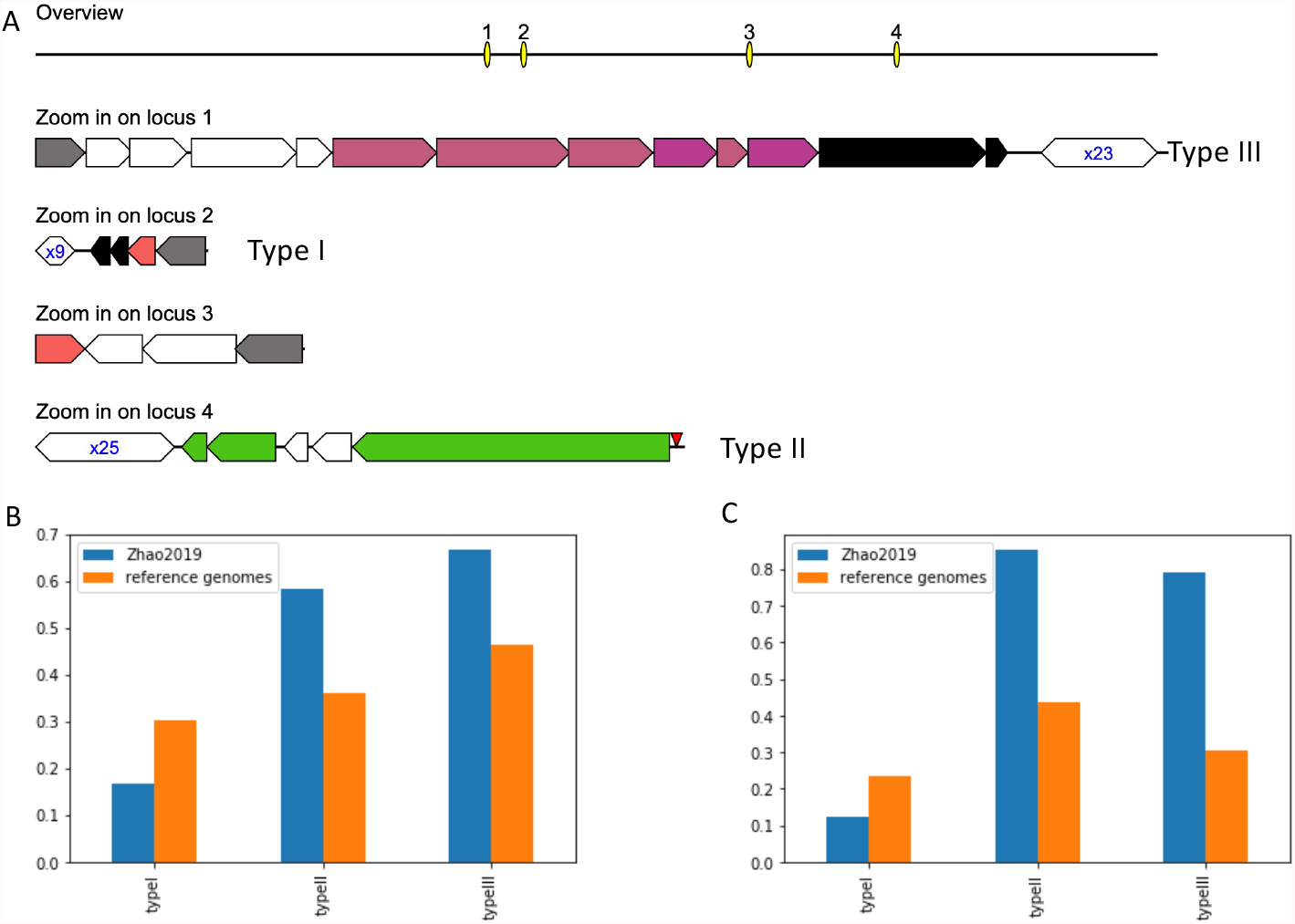
The three types of CRISPR-Cas systems found in *B. fragilis* genomes. (A) Representatives of *B. fragilis* CRISPR-Cas systems found in *B. fragilis* strain S14 (accession number: GCA 001682215.1 ASM168221v1). (B) Prevalence of CRISPR-Cas systems among the NCBI reference genomes and Zhao2019 isolates. (C) Spacer content heterogeneity of the CRISPR arrays.

**Table 1:**
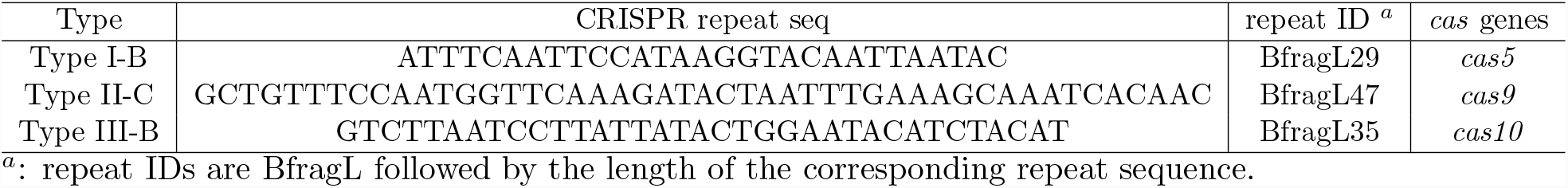
Signature *cas* genes and CRISPR repeat sequences for the three types of CRISPR-Cas systems found in *B. fragilis*

### Inter-subject spacer diversity of *B. fragilis* CRISPR-Cas systems

An evaluation of the CRISPR-Cas system distribution among all isolates of the Zhao2019 dataset showed that CRISPR-Cas system types were unevenly and dis-similarly distributed between individuals (Figure 2B and Table 2). Type I-B CRISPR-Cas systems were among the least prevalent with only isolates from two individuals (S07 and S08) containing this type of CRISPR-Cas system. Type II-C and Type III-B CRISPR-Cas systems are similarly prevalent with isolates from six individuals (see Table 2 for the lists of individuals that contain these systems). No CRISPR-Cas systems were found to be present within Zhao2019 Subjects S04, S05, and S12. The lack of uniformity of CRISPR-Cas system presence, or lack thereof, suggests that lineages of *B. fragilis* between individuals are, for the most part, unique from each other, reaffirming the findings of Zhao et al. Similarly, there was an observed lack of shared inter-individual spacer content, with majority of the spacers observed being individual specific (Figure 4), with the most common shared spacer being the anchor spacer on the trailer end of observed CRISPR arrays.

**Table 2:** Presence of the different types of CRISPR-Cas systems in the 12 individuals.

| Type | S01 (t10) | S02 (t2) | S03 (t2) | S04 (t2) | S05 (t3) | S06 (t2) | S07 (t3) | S08 | S09 | S10 | S11 | S12 |
| --- | --- | --- | --- | --- | --- | --- | --- | --- | --- | --- | --- | --- |
| Type I-B |  |  |  |  |  |  | yes | yes |  |  |  |  |
| Type II-C |  | yes | yes |  |  | yes | yes |  |  | yes |  |  |
| Type III-B | yes | yes |  |  |  | yes | yes |  | yes | yes |  |  |

Spacer content heterogeneity score (Figure 2) and compressed spacer graph of Type I-B CRISPR-Cas systems (Figure 3A) found within Zhao2019 isolates shows that the CRISPR array structure of Type I-B systems have low heterogeneity and are less active in terms of spacer turnover compared to other CRISPR-Cas systems in *B. fragilis*. In comparison, Type III-B CRISPR-Cas Systems contained mostly individual specific spacers and shared very few spacers between individuals. This pattern of individual-specific spacers is reflected in the branching structures observed in the compressed spacer graphs (Figure 3B). Each branch within the spacer graph represents an unique CRISPR array structure; bottle-neck nodes (e.g. 9, 35, 40) represent uniformly shared spacer(s) in spacer sharing CRISPR arrays. The observed branching structure in the compressed spacer graph indicates a diverse CRISPR array structure between individuals, indicative of the activity and heterogeneity of Type III-B CRISPR-Cas systems within *B. fragilis*. For comparison, the spacer content hetegenerity score shows similar trends among the *B. fragilis* reference genomes (see Figure 2C).

**Figure 3:**
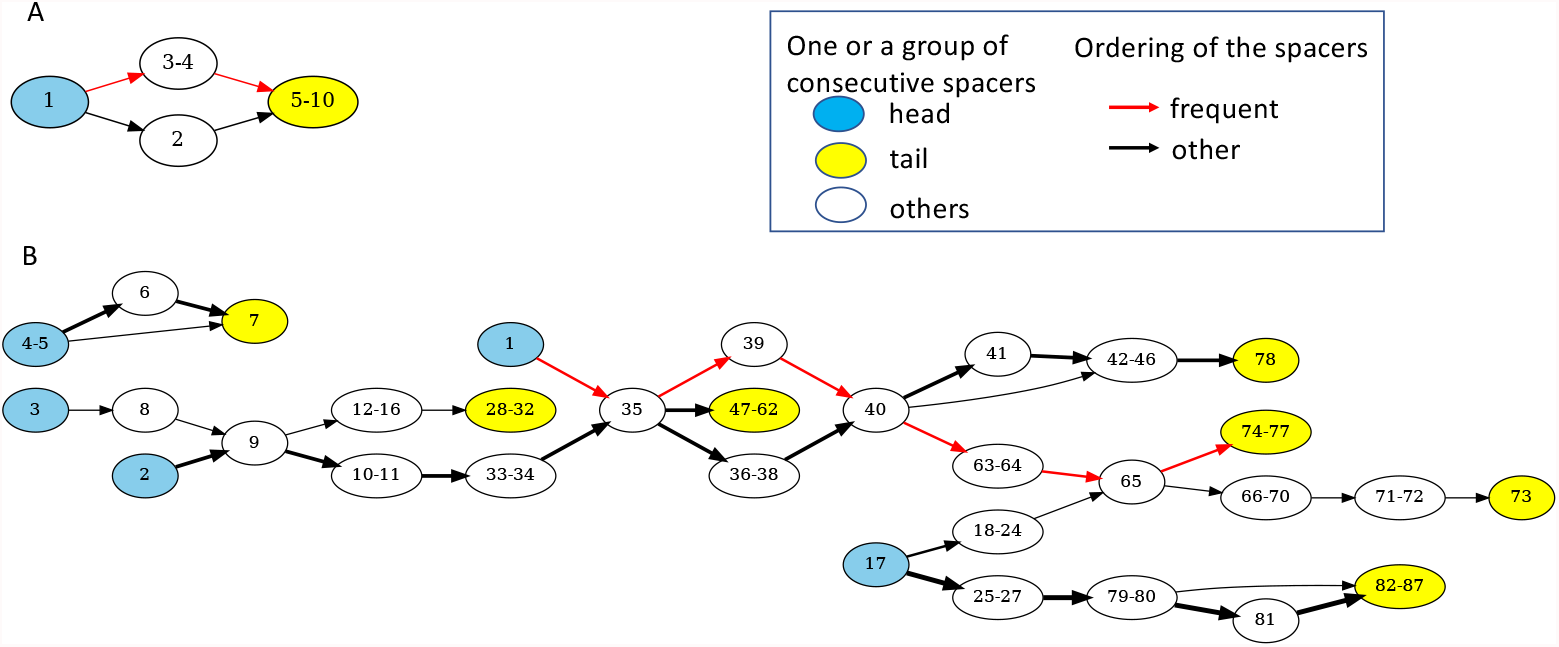
Compressed spacer graphs of Type I-B (A) and Type III-B (B) CRISPR-Cas systems in *B. fragilis*. Nodes represent a single or several consecutive spacers, and edges represent the ordering of the spacers in arrays. Nodes containing leader end spacers are highlighted in blue, and nodes containing trailer end spacers are shown in yellow.

The Type II-C CRISPR-Cas systems found within the Zhao2019 isolates were among the most diverse between the three observed types of CRISPR-Cas systems found within *B. fragilis*. Among the seven subjects that contained Type II-C CRISPR-Cas systems, many of the identified Type II-C CRISPRs did not share inter-subject spacers, except for trailer spacers (i.e., end spacers; one example is node #112 shown in Figure 4) which have been previously hypothesized as ancient spacers or anchor spacers [58, 30, 47]. The spacer sequence diversity can be seen in Figure 4, where each branch path represents a unique CRISPR array observed. The diversity of the CRISPR arrays observed in Type II-C CRISPR-Cas systems suggests that Type II-C systems have greater spacer activity (e.g. spacer acquisition and loss), and also highlight the evolutionary pressures that MGEs exert on *B. fragilis*.

**Figure 4:**
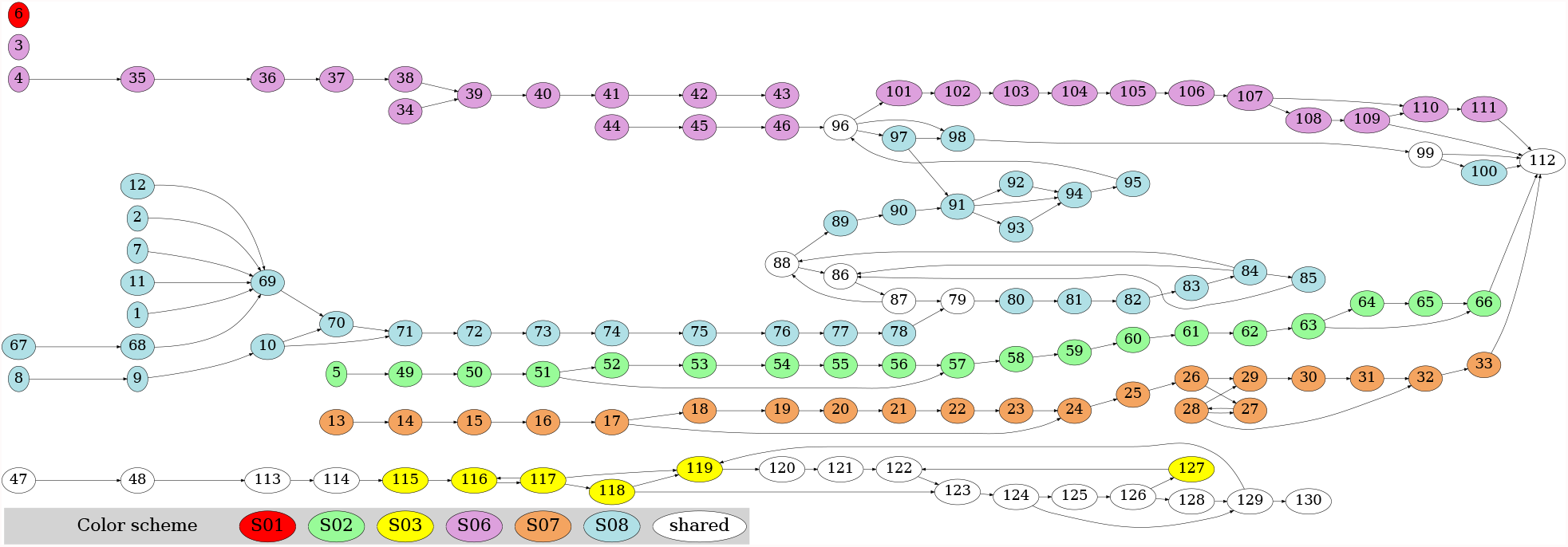
Spacer graph of the Type II-C CRISPR-Cas systems in *B. fragilis*. Each node represents a unique spacer. Spacers that are unique to individuals are highlighted in different colors (see color scheme in the plot; spacers shared by isolates from at least two individuals are shown in white).

### Intra-individual CRISPR-Cas dynamics of *B. fragilis*

Taking advantage of the temporal intra-individual isolates of *B. fragilis*, we were able to study micro-dynamics of *B. fragilis* community dynamics and the adaptation of its CRISPR arrays over time per individual. Overall CRISPR array structures remain relatively stable across samples from the same individual, with some slight variations between observed CRISPR arrays (e.g Figure 4 S01, S02, S03, S06, S07). A notable example is the arrays of type III-B CRISPR-Cas systems found in S01 isolates (Figure 5A). Isolates were derived from this individual at 10 different time points spanning more than 2 years, and we only observed a small variation of the arrays in those isolates, resulting in a simple spacer graph with only one branching structure involving the loss (or gain) of a spacer in the middle of the arrays. However, in some instances, periods of diverse spacer acquisition were observed from samples from the same individual (e.g., S08). As shown in Figure 4, various strains of *B. fragilis* with varying CRISPR array structures were observed from isolates obtained in the same individual (S08) at a single time point. The spacer graph shows a funneling pattern, where multiple nodes on the leader end of the CRISPR array converge into a single neighboring node (see blue nodes on the left of Figure 4). This observed pattern in the spacer graph suggests that multiple “lineages” have gained alternative leader end spacers in comparison to each other, specifically when the bacteria are exposed to different MGEs and are evolving according to the observed threat. As another example, there are multiple “lineages” of *B. fragilis* containing different Type II-C CRISPR arrays in S02 at time point 1 (S02-0001), one of which was also present in S02 at a later time point (S02-0024, 24 days apart from the first time point) (see Figure 5B). Similarly, there are also multiple lineages of *B. fragilis* containing diverse Type II-C CRISPR arrays in S06 (S06-0001) (see Figure 5C). While previous studies [59, 60, 44] have shown that intra-individual populations of *B. fragilis* are dominated by a single strain, our findings here show that in some cases many “lineages”, or strains of *B. fragilis*, remain present within the same individual at any given time point. The observation of various intra-individual *B*.*fragilis* strains is yet another example of the evolutionary arms race between host and the invading MGE.

**Figure 5:**
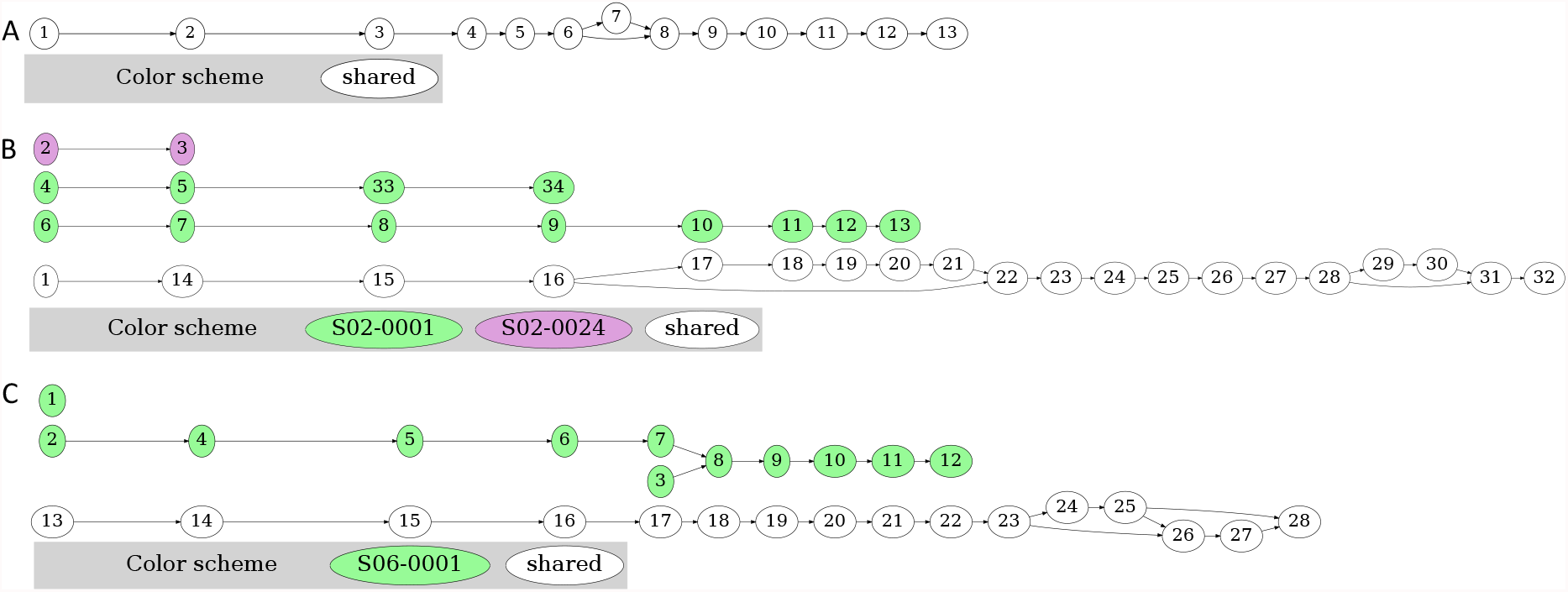
Intra-individual CRISPR array diversity. (A) Spacer graph of CRISPR arrays from individual S01 isolates, consisting of 10 time points, showing low spacer diversity. (B) Spacer graph for individual S02. Green nodes are unique spacers identified from time point 1 (S02-0001), purple nodes are from time point 24 (S02-0024), and white nodes represent spacers found in multiple time points. (C) Spacer graph for individual S06.

### Interaction network of *B. fragilis* and its invaders

Among the 1531 unique spacers identified from *B. fragilis* isolates, 522 found matches (protospacers) in 161 MGEs (153 phages and 8 plasmids). 108 out of the 153 phages could be assigned to a family by PhaGCN with a majority of them being Siphoviridae (93, 86%). Using these spacers, interaction networks between *B. fragilis* and its invaders were inferred.

Analysis of the networks (Figure 6A and B) showed varying levels of micro-dynamics within *B. fragilis* CRISPR-Cas systems. The spacer-MGE network (Figure 6A) contains a few modules each containing a large number of MGEs and spacers (e.g., modules a, b, c and d highlighted in the Figure), likely a result of the arms-race between *B. fragilis* and MGEs (*B. fragilis* acquired new spaces to maintain immunity and invaders mutated to evade immunity). The spacer-MGE network shows that *B. fragilis* used its Type I-B and II-C CRISPR-Cas systems extensively to defend against MGEs that were mostly phages (the network contains 353, 163, and 3 spacers that were exclusively caught in Type II-C, Type I-B, and Type III-B CRISPR-Cas systems, respectively). It also suggests differential defense activities of the Type I-B and II-C CRISPR-Cas systems against some invaders (e.g., those included in modules a and b were preferentially targeted by Type II-C CRISPR-Cas systems; by contrast, invaders included in modules c and d don’t show such preference). Figure 6B (focusing on *B. fragilis* isolates from several individuals) showed that some invaders (such as P1, P2, P3 and P4 located at the center of the network) have their traces found in *B. fragilis* in many different individuals, likely the result of ubiquitous presence of these MGEs in human gut. Despite of these central MGEs that make the whole network highly connected, we observed groupings of *B. fragilis* isolates from one or two subjects with some more localized MGEs (e.g., the MGEs that were targeted by the *B. fragilis*’ CRISPR-cas systems in individual S09). Figure 6C shows the distribution of protospacers in NC 011222 (Bacteroides phage B40-8, labelled as P4 in Figure 6B) that was targeted extensively by both Type I-B and Type II-C CRISPR-Cas systems.

**Figure 6:**
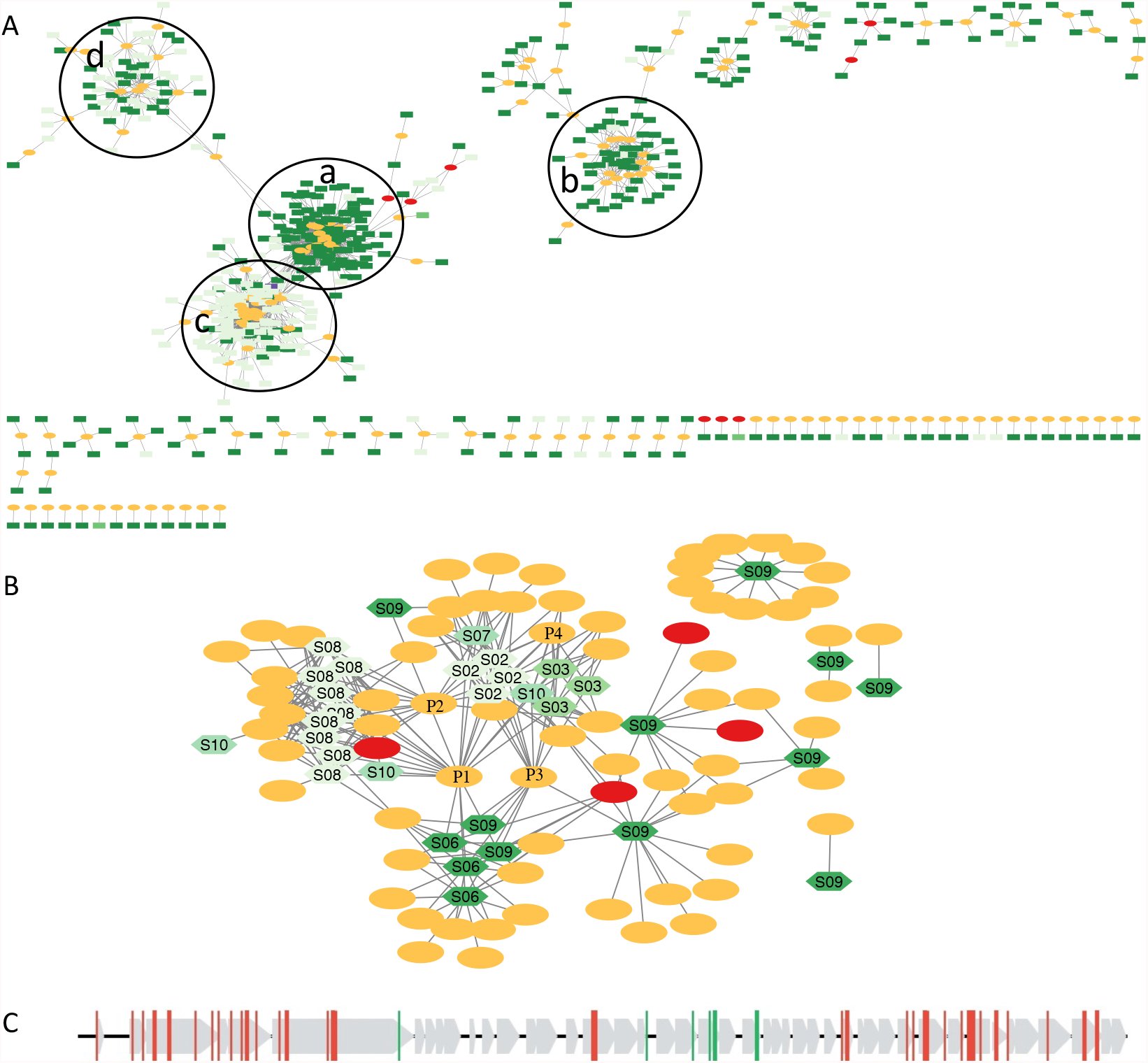
Interactions between *B. fragilis* and MGEs shown as a spacer-MGE (A) and host-MGE network (B). (A) includes spacers identified from Zhao2019 and the reference genomes that had their sources (i.e., protospacers) identified; by contrast, (B) includes spacer-contributing *B. fragilis* isolates from Zhao2019 only. MGEs containing protospacers are shown in ovals with plasmids highlighted in red and phages in orange. In (A), spacers identified in different types of CRISPR-Cas systems are shown in rectangles of different colors (Type I-B in light green, Type II-C in dark green, and Type III-B in blue). In (B), hosts are shown in hexagons with *B. fragilis* isolates from different individuals shown in different shades of green; four phage MGEs are highlighted with labels: P1 (uvig 425872, Siphoriridae), P2 (uvig 422350, Siphoviridae), P3 (k141 68 round8-12 1718861, Microviridae), and P4 (NC 011222, Siphoriridae). (C) shows the distribution of protospacers in NC 011222, with green and red vertical lines representing protospacers matching spacers found in Type I-B and Type II-C CRISPR-Cas systems, respectively (gray arrows represent the genes).

## Discussion

In this paper, we expanded upon previous works [39] and explored the CRISPR-Cas dynamics within *B. fragilis* genomes, while focusing on dynamics pertaining to a time-resolved study of *B. fragilis* within and between individuals. We analyzed a total of 823 genomes, a 7.5 fold difference in number of genomes analyzed in previous *B. fragilis* CRISPR-Cas papers [39]. While *B. fragilis* is a common commensal bacterium of the human gut microbiome, sometimes a probiotic candidate and sometimes pathogen, its role as one of the most virulent members of the *Bacteroides* genus should not be overlooked [61]. Part of *B. fragilis* virulence is due to its potent virulence factors, and as such, a thorough understanding of the mechanisms and factors that contribute to its virulence, horizontal gene transfer, and evolution are important. By utilizing CRISPR-Cas systems and focusing on time series isolates, we were able to reveal micro-dynamics found in *B. fragilis* isolates within and between individuals.

The analysis of NCBI’s reference genomes and genomes from the Zhao2019 dataset enabled us to update the evaluation of known CRISPR-Cas systems found within *B. fragilis*. Particularly, we found three types of CRISPR-Cas systems (Type I-B, Type II-C, and Type III-B) with varying distributions among the genomes. Our analysis also shows that a fourth previously reported CRISPR-Cas system in *B. fragilis* was a false CRISPR-like artifact. This CRISPR-like artifact was previously characterized as an orphaned CRISPR array [39], but due to its structure containing only two spacers, three repeats, as well as non-uniform repeat sequences, we believe this is not an orphaned CRISPR array.

By focusing our analysis on bacterial isolates, we were also able to reduce the chance of artifacts (e.g., chimeric contigs) from the assembly process, thereby reducing the noise in our analysis of the evolutionary processes in *B. fragilis* CRISPR-Cas systems. Our analysis shows that while all *B. fragilis* CRISPR-Cas system types had some level of plasticity, where CRISPR arrays across different time points and individuals were heterogeneous, the level of heterogeneity varied between CRISPR-types and even time-points. Intra-individual variations of CRISPR arrays, such as those found in Individual S08 (Figure 4), showed periods of rapid expansion and diversification of CRISPR spacers between strains of observed isolates; these periods of diversification can be observed in the branching structures of the spacer graph. In comparison, periods of contraction where little to no CRISPR spacer content heterogeneity was observed were similarly present in intra-individual CRISPR-Cas systems, such as those found in Individual S01 (Figure 5A). Unsurprisingly, most inter-individual CRISPR-Cas systems did not share many spacers between individuals. This suggests that strains found within individuals were derived from a common ancestor, but strains have since diverged (as evident in their spacer content and exposure to different MGEs) and have not been circulating between individuals. Here we also show that CRISPRs can go through periods of expansion, while others go through periods of stability, suggesting that CRISPR evolution is not a constant process but occurs in modes. Uncovering these CRISPR-Cas dynamics would not be possible without time series analysis of the same bacterial lineage.

We found that *B. fragilis* CRISPR-Cas systems seemed to prefer targeting phage genomes over plasmid genomes while exploring the interplay/dynamics of *B. fragilis* and its MGEs. This is a contrast to some studies which found CRISPRs favoring the targeting of plasmids over phages [62, 63]. CRISPR spacer-MGE networks also revealed micro-dynamics of *B. fragilis* CRISPR targets, where we observed several notable network structures. Hairball-like structures, where a single spacer targeted many unique MGE targets, and exemplified that in some cases CRISPR spacers were likely able to target multiple MGEs through the same CRISPR spacer. This suggests that the protospacer is conserved across many targets. In addition to hairball like structures, it was also observed that several spacer nodes and MGE nodes formed cliques/modules, where nodes clustered together more closely to each other than other members of the network. Within these modules, MGE nodes shared an edge with many spacer nodes, suggesting that these MGEs contained many protospacers. This observation of many spacers targeting the same MGE may be suggestive of a process known as ‘primed CRISPR adaptation’. In primed CRISPR adaptation, the presence of an existing spacer is used to enhance the acquisition of new spacers on the same MGE target [64, 65]. Alternatively, it may be possible that these instances of multiple targeting are a result of naive adaptation where spacers were independently acquired.

Not all spacers identified in *B. fragilis* had a matching MGE protospacer target, which might have biased our analysis to spacer targets based on available MGE database genomes. However, it has been suggested that most unidentified spacers relate to host-specific mobile elements [66, 67] and thus without adequate sequencing and annotation of the hosts’ microbiome, many of the spacer targets will remain unresolved. Another hypothesis to the limited spacer-MGE associated matches, especially in trailer end (older) spacers, is that protospacer sites of targeted MGEs have since mutated to evade detection by the CRISPR spacer and the MGE target pre-dates sequencing technology; thus, spacers are unable to match to any known protospacer targets within the available MGE databases.

Additionally, in compressed spacer graphs, we observed periods of expansion and contraction of CRISPR arrays. Funneling patterns are of particular interest and were mostly observed at the leader end of spacer graphs. The lack of these funnel shaped patterns in the middle or trailer end of compressed spacer graphs suggests that certain spacers may provide an evolutionary advantage compared to other spacers, and establish itself as the dominant strain, out competing strains containing less fit CRISPR arrays; thus we do not see this branching structure in ‘older’ areas of the CRISPR array.

By exploring CRISPR-Cas systems present in *B. fragilis* and the dynamics of its host-MGE networks, we uncovered micro-dynamics of *B. fragilis* adaptation against invaders. While our work improves the understanding of *B. fragilis* adaptation to MGE exposure, more work is still needed to understand how CRISPR adaptation plays a role in *B. fragilis* acquisition of virulence factors, evolution, and horizontal gene transfer. In particular, one main challenge to Host-Invader analysis is the limitation of available MGE databases. Future efforts and resources to maintain databases of MGEs and other elements of the microbiome (e.g. fungome) remain invaluable for further understanding of the microbiome, and not just prokaryotic members. A better understanding of how *B. fragilis* and other pathobionts interact with their invading mobile elements will enable a better understanding of their evolution and the elements responsible for their pathogenicity.

## Availability of the results

We made the CRISPR-one annotations of all the genomes available for download along with visualization as a web resource at https://omics.informatics.indiana.edu/CRISPRone/Bfragilis. Phages and plasmids that were predicted to be targeted by the CRISPR-cas systems, and their interaction networks are also available at the web resource.

